# Clinical Characterization of Host Response to Simian Hemorrhagic Fever Virus Infection in Permissive and Refractory Hosts: A Model for Determining Mechanisms of VHF Pathogenesis

**DOI:** 10.1101/454462

**Authors:** Joseph P. Cornish, Ian N. Moore, Donna L. Perry, Abigail Lara, Mahnaz Minai, Dominique Promeneur, Katie R. Hagen, Kimmo Virtaneva, Monica Paneru, Connor Buechler, David H. O’Connor, Adam L. Bailey, Kurt Cooper, Steven Mazur, John G. Bernbaum, James Pettitt, Peter B. Jahrling, Jens H. Kuhn, Reed F. Johnson

**Affiliations:** Emerging Viral Pathogens Section, Laboratory of Immunoregulation, Division of Intramural Research, National Institute of Allergy and Infectious Diseases, National Institutes of Health, Frederick, Maryland, USA; Infectious Disease Pathogenesis Section, Comparative Medicine Branch, Division of Intramural Research, National Institute of Allergy and Infectious Diseases, National Institutes of Health, Rockville, Maryland, USA; Integrated Research Facility, Division of Clinical Research, National Institute of Allergy and Infectious Diseases, National Institutes of Health, Frederick, Maryland, USA; Genomics Unit, Research Technologies Branch, Rocky Mountain Laboratories, National Institute of Allergy and Infectious Diseases, National Institutes of Health, Hamilton, Montana, USA; Department of Pathology and Laboratory Medicine, University of Wisconsin–Madison, Madison, Wisconsin 53711, USA; Wisconsin National Primate Research Center, Madison, Wisconsin 53711, USA

**Author notes:** Current Affiliation: Department of Pathology and Immunology, Washington University in St. Louis School of Medicine, St. Louis, MO, 63110, USA. Address correspondence to Reed F. Johnson. Present address: One MedImmune Way, Gaithersburg, Maryland, USA.

**Keywords:** *Arteriviridae*, arterivirus, host-response, macaque, patas monkey, pathogenesis, SHFV, simartevirus, simian hemorrhagic fever, VHF, viral hemorrhagic fever

## Abstract

Simian hemorrhagic fever virus (SHFV) causes a fulminant and typically lethal viral hemorrhagic fever (VHF) in macaques (Cercopithecinae: *Macaca* spp.) but causes subclinical infections in patas monkeys (Cercopithecinae: *Erythrocebus patas*). This difference in disease course offers a unique opportunity to compare host-responses to infection by a VHF-causing virus in biologically similar susceptible and refractory animals. Patas and rhesus monkeys were inoculated side-by-side with SHFV. In contrast to the severe disease observed in rhesus monkeys, patas monkeys developed a limited clinical disease characterized by changes in complete blood counts, serum chemistries, and development of lymphadenopathy. Viremia was measurable 2 days after exposure and its duration varied by species. Infectious virus was detected in terminal tissues of both patas and rhesus monkeys. Varying degrees of overlap in changes in serum concentrations of IFN-γ, MCP-1, and IL-6 were observed between patas and rhesus monkeys, suggesting the presence of common and species-specific cytokine responses to infection. Similarly, quantitative immunohistochemistry of terminal livers and whole blood flow cytometry revealed varying degrees of overlap in changes in macrophages, natural killer cells, and T-cells. The unexpected degree of overlap in host-response suggests that relatively small subsets of a host’s response to infection may be responsible for driving pathogenesis that results in a hemorrhagic fever. Furthermore, comparative SHFV infection in patas and rhesus monkeys offers an experimental model to characterize host-response mechanisms associated with viral hemorrhagic fever and evaluate pan-viral hemorrhagic fever countermeasures.

**IMPORTANCE:** Host-response mechanisms involved in pathogenesis of VHFs remain poorly understood. An underlying challenge is separating beneficial, inconsequential, and detrimental host-responses during infection. The comparison of host-responses to infection with the same virus in biologically similar animals that have drastically different disease manifestations allows for the identification of pathogenic mechanisms. SHFV, a surrogate virus for human VHF-causing viruses likely causes subclinical infection in African monkeys such as patas monkeys but can cause severe disease in Asian macaque monkeys. Data from the accompanying article by Buechler *et al*. support that infection of macaques and baboons with non-SHFV simarteviruses can establish persistent or long-term subclinical infections. Baboons, macaques, and patas monkeys are relatively closely taxonomically related (Cercopithecidae: Cercopithecinae) and therefore offer a unique opportunity to dissect how host-response differences determine disease outcome in VHFs.

## INTRODUCTION

Viral hemorrhagic fevers (VHFs) are primarily caused by single-stranded RNA viruses (<u>1</u>). VHF is a broadly defined syndrome: fever, hepatic and renal complications, large increases in proinflammatory cytokines and coagulopathy are common features (<u>2</u>, <u>3</u>). Simian hemorrhagic fever virus (SHFV) is a positive-sense, single-stranded RNA virus classified in the family *Arteriviridae*, which also includes equine arteritis virus and porcine reproductive and respiratory syndrome viruses 1 and 2 (<u>4</u>, <u>5</u>). In addition to SHFV, several other simian arteriviruses (genus *Simartevirus*) have been identified (<u>6</u>-<u>8</u>). Among simarteviruses, SHFV, simian hemorrhagic encephalitis virus (SHEV), and Pebjah virus (PBJV) are known to cause severe disease in Asian macaques of various species (<u>9</u>). Kibale red colobus virus 1 (KRCV-1) was found to cause a self-limiting disease in crab-eating macaques (<u>10</u>). It is not known whether the other identified simarteviruses infect or cause disease in macaques or their natural hosts. Here, and in combination with accompanying article by Buechler *et al*., we examine infection of natural hosts (patas monkeys and baboons) with simarteviruses, and compare disease course of these simarteviruses in macaque monkeys.

SHFV was discovered during a VHF outbreak in National Institutes of Health (NIH) animal facilities in the United States in 1964 (<u>11</u>, <u>12</u>). Transmission during the NIH SHFV outbreak is thought to have occurred through tattooing needles used on both macaques and co-housed African primates (<u>12</u>). The virus is highly virulent in rhesus monkeys (*Macaca mulatta*), crab-eating macaques (*Macaca fasicularis*), stump-tailed macaques (*Macaca arctoides*), and Japanese macaques (*Macaca fuscata*), but SHFV causes little to no disease in African primates such as patas monkeys or baboons (<u>11</u>, <u>13</u>, <u>14</u>). SHFV infection in macaques mirrors aspects of human VHFs, such as Ebola virus disease, by inducing fever, edema, coagulopathy, hepatocellular degeneration and necrosis, and elevated inflammatory cytokine concentrations (<u>13–15</u>). Like all VHFs, simian hemorrhagic fever (SHF) is thought to be driven by a dysregulated host-response leading to a dysregulated immune response and poor viral clearance (<u>13</u>, <u>14</u>, <u>16</u>).

Studying SHFV offers a unique opportunity to compare infection and associated responses in refractory and highly susceptible primates that are biologically similar to each other and to humans. Some hemorrhagic fever-causing viruses naturally infect non-primate mammals that may serve as hosts or reservoirs. For instance, Marburg virus (*Filoviridae: Marburgvirus*, MARV) and Ravn virus (*Filoviridae: Marburgvirus*, RAVV) naturally circulate in Egyptian rousettes (*Rousettus aegyptiacus*), in which they do not cause disease, whereas these viruses cause frequently lethal disease experimentally in primates and naturally in humans (<u>17</u>). Similarly, arenaviruses associated with human VHFs, such as Machupo virus and Lassa virus (*Arenaviridae: Mammarenavirus*), subclinically infect distinct rodent reservoir hosts (<u>18</u>, <u>19</u>).

Comparing the response to infection between refractory hosts, preferably the reservoirs themselves, and susceptible hosts may offer significant insight into responses involved in VHF pathogenesis. For most hemorrhagic fever-causing viruses, comparisons between refractory and susceptible animals during infection is confounded by large biological differences. For example, work with pathogenic mammarenaviruses has demonstrated that mechanisms present in murine hosts, even susceptible ones, are fundamentally different than those of primates (<u>20</u>). Unlike these examples, SHFV infects biologically similar refractory and susceptible animals and may provide a path towards meaningful interspecies comparisons of responses to hemorrhagic fever-causing virus infection.

The goal of this work was to compare host-responses in biologically similar nonhuman primates, patas (refractory) monkeys and rhesus (susceptible) monkeys, infected with SHFV to identify factors associated with differential outcomes to infection. The findings are the first in-depth characterization of SHFV infection in patas monkeys and confirm that patas monkeys are largely unaffected by SHFV infection. Our work demonstrates that, although patas and rhesus monkeys develop drastically different diseases, the host-responses to infection overlap, and suggest that VHF pathogenesis may be driven by a relatively small subset of the overall host-response to infection.

## RESULTS

### Simian hemorrhagic fever virus (SHFV) infection results in mild clinical disease

Twelve monkeys were grouped by species into SHFV-inoculated and PBS inoculated groups, n=3 per. Inoculations were 1-ml intramuscular injections in the right quadriceps. The 3 SHFV-inoculated patas monkeys developed axillary and inguinal lymphadenopathy starting on day 10 post-inoculation (PI) that persisted until the conclusion of the experiment at 21 days PI. The 3 SHFV-inoculated rhesus monkeys developed severe disease. Two subjects (“non-survivors”) met clinical endpoint criteria and were euthanized on days 8 and 11 post-inoculation. The third subject (“survivor”) survived until the conclusion of the experiment (day 20 PE). Signs of disease were first detectable on day 4 PI SHFV-inoculated rhesus monkeys developed a range of clinical signs and included: gingival bleeding (1/3 subjects), inguinal lymphadenopathy (1/3), and axillary lymphadenopathy (2/3). All SHFV-inoculated rhesus monkeys developed petechial rashes and axial and inguinal lymphadenopathy by their respective endpoints. All 3 SHFV-inoculated rhesus monkeys developed tremors and motor dysfunction by day 6 PI that remained until each subject’s respective endpoint. The non-surviving rhesus monkeys developed facial edema that began on day 8 PI that persisted to their respective endpoints. The surviving SHFV-inoculated rhesus monkey developed facial edema that began on day 12 PI and resolved by day 16 PI. All mock-inoculated rhesus appeared clinically normal and displayed no outward signs of disease throughout the experiment.

### SHFV infection results in clinical pathology changes in patas monkeys

In SHFV-inoculated patas monkeys, alanine aminotransferase (ALT), alkaline phosphatase (ALP) and aspartate aminotransferase (AST) concentrations were elevated on day 4 PI and remained above baseline until the end of the experiment (Fig 1A-C). γ-glutamyl transferase (GGT) concentrations remained normal in all SHFV-inoculated patas monkeys (Fig 1D). SHFV-inoculated rhesus monkeys showed similar trends except that GGT concentrations rose starting on day 4 PI. ALP, AST, and GGT concentrations remained elevated in the surviving SHFV-inoculated rhesus monkey until the conclusion of the experiment, whereas ALT returned to baseline concentrations. Although changes in serum chemistries were observed in all SHFV-inoculated subjects, values did not reach concentrations suggestive of a severe clinical disease. All subjects experienced decreases in albumin concentration coinciding with globulin concentration increases (Fig 1F) on day 8 and day 4 PI in SHFV-inoculated patas and rhesus monkeys, respectively. Reticulocyte counts decreased in both SHFV-inoculated patas and rhesus monkeys: on day 6 PI in patas monkeys and on day 4 PI in rhesus monkeys (Fig 1E), but did not drop below the normal range. Hematocrit (HCT) remained normal in all subjects except for the surviving SHFV-inoculated rhesus where HCT decreased starting on day 10 PI with anemia persisting to study end (Fig 1G). Albumin concentrations (Fig 1H) began decreasing in SHFV inoculated patas monkeys on day 6 PI to day 8 PI after which concentrations remained depressed till the conclusion of the study. In SHFV-inoculated rhesus monkeys, albumin concentrations began decreasing on day 2 and continued to decrease in all subjects till their respective endpoints. No significant changes in serum chemistry were observed in mock-inoculated patas and rhesus monkeys.

**Figure 1.**
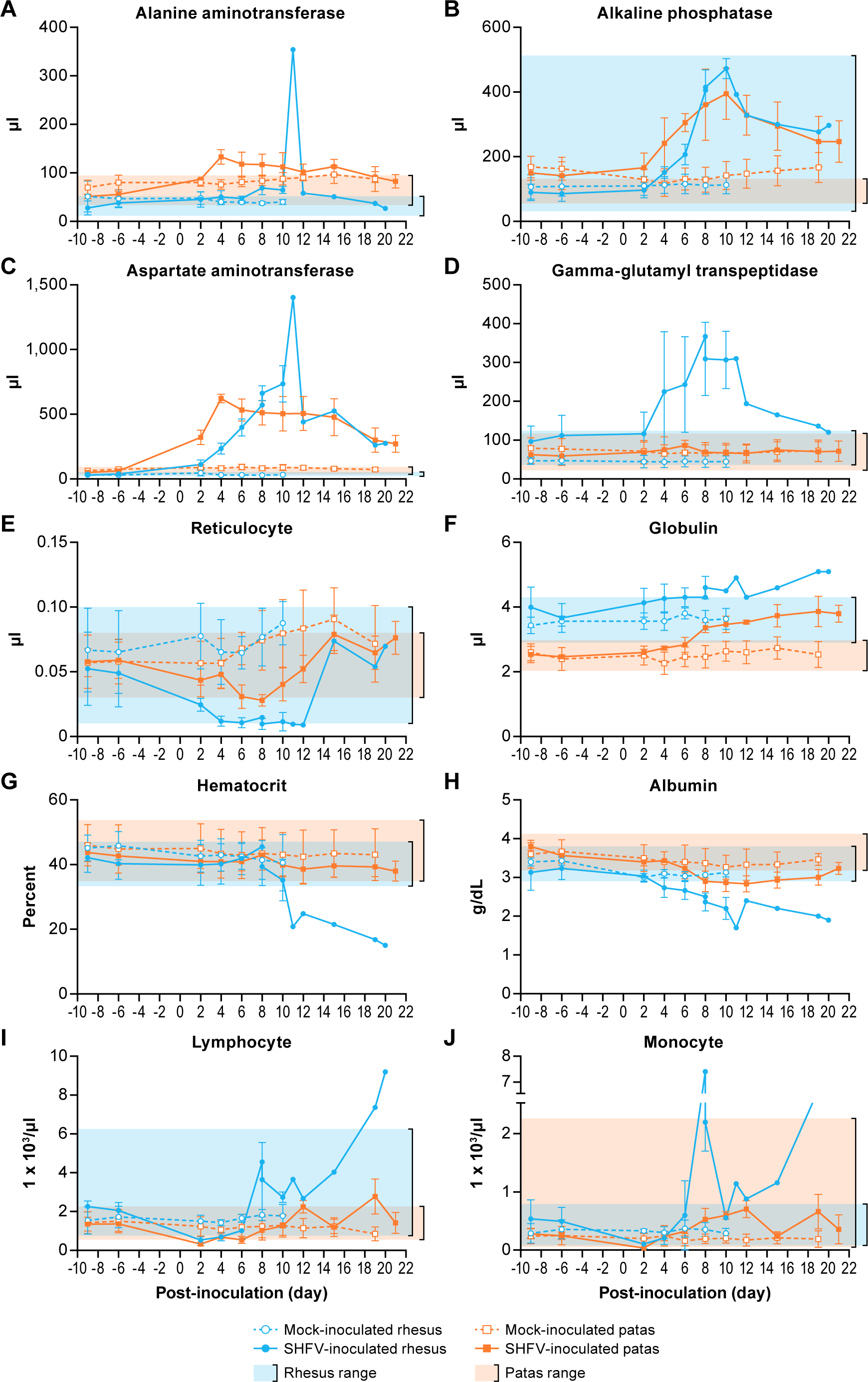
Alanine aminotransferase (A), alkaline phosphatase (B), aspartate aminotransferase (C), gamma-glutamyltransferase (D), reticulocyte number (E), globulin (F), hematocrit (HCT) (G), albumin (H), lymphocyte number (I), monocyte number (J) values for patas monkeys (orange lines) and rhesus monkeys (blue lines) either inoculated with 5,000 PFU of SHFV (open symbols) or with PBS (closed symbols). Shaded regions represent standard range of all pre-exposure values for patas monkeys or previously collected values for rhesus monkeys. Data represent means of each group.

Complete blood counts revealed minor decreases in lymphocyte numbers early in all SHFV-inoculated patas and rhesus monkeys, followed by a return to baseline counts at day 8 PI (Fig 1I). Lymphocyte counts increased in the surviving rhesus, but the mechanism is not known at this time and may represent further maturation of the immune response. Monocyte counts decreased slightly in all SHFV-inoculated patas and rhesus monkeys on day 2 PI before increasing by day 6 PI. After day 6 PI monocyte counts (Fig 1J) in SHFV-inoculated patas monkeys returned to baseline by day 15 followed by a second increase in 2/3 subjects. Unlike the patas monkeys, monocyte counts in SHFV-inoculated rhesus monkeys continued to increase until each subject’s endpoint. No significant changes were observed in complete blood counts in any mock-inoculated subjects.

### SHFV infection causes mild pathology in patas monkeys

Gross examination of SHFV-inoculated patas monkeys during necropsy confirmed peripheral lymphadenopathy in all three subjects, but there were no other remarkable findings. Non-surviving SHFV-inoculated rhesus monkeys had marked hepatosplenomegaly: hepatic tissues were friable and firm. Moderate peripheral and visceral lymphadenopathy was found in both non-surviving rhesus monkeys, whereas moderate peripheral lymphadenopathy was the only significant finding in the surviving rhesus monkey. The kidneys of one non-surviving SHFV-inoculated rhesus monkey contained multiple infarctions with severe renal hemorrhage and necrosis. No significant gross lesions were observed in either mock-inoculated group.

Histopathological examination of the spleen in SHFV-inoculated patas monkeys revealed abundant plasma cells in one subject. The spleens of the 2 remaining SHFV-inoculated patas monkeys were within normal limits. The livers of two SHFV-inoculated patas monkeys had minimal inflammatory changes, whereas that of the third patas monkey was within normal limits. Hyperplasia was evident in the inguinal lymph nodes of 1 SHFV-inoculated patas monkey. The spleens of SHFV-inoculated rhesus monkeys were different in each subject: in the two non-survivors, one was congested, whereas fibrin deposition was evident in the other. The spleen of the surviving subject exhibited changes that were consistent with reactive lymphoid hyperplasia, characterized by diffuse expansion and proliferation of B-cells at the margins of each follicle. Each of the livers of SHFV-inoculated rhesus monkeys were also histologically different. In the survivor, perivascular inflammation with multifocal areas of necrosis were evident. The major finding in the liver of one non-survivor was vacuolated hepatocytes, whereas rare thrombi were the major observation in the remaining non-survivor. Hyperplasia was present in inguinal lymph nodes of all three SHFV-inoculated rhesus monkeys. Tissues of all mock-inoculated subjects were normal apart from vascular congestion in 2 patas monkey spleens. While all three SHFV-inoculated rhesus monkeys displayed neurological signs, on histopathological examination CNS tissues did not reveal any evidence of vasculitis or other changes that would suggest encephalitis. CNS tissues of all mock-inoculated animals and SHFV-inoculated patas monkeys were found to be within normal histologic limits. Upon histopathological examination kidneys of all subjects appeared to be within normal histologic limits.

Immunohistochemical staining used to detect ionized calcium-binding adaptor 1 (Iba-1)-expressing macrophages revealed morphologically-normal macrophages in the livers of all 3 SHFV-inoculated patas monkeys. In contrast, the livers of all 3 SHFV-inoculated rhesus monkeys contained macrophages that were often rounded and contained a diffusely vacuolated cytoplasm (Fig 2A, inset). These changes in macrophage morphology were in direct contrast to these cells in infected patas monkeys which often exhibited a more stellate shape and cytoplasm that was diffusely dark brown when evaluated immunohistochemically. Similar findings were seen in the inguinal lymph nodes and spleen of SHFV-inoculated patas and rhesus monkeys, although rounded macrophages were detected in the spleen of one SHFV-inoculated patas monkey and inguinal lymph nodes of a second SHFV-inoculated patas monkey. Macrophages appeared morphologically normal in all mock-inoculated subjects.

**Figure 2.**
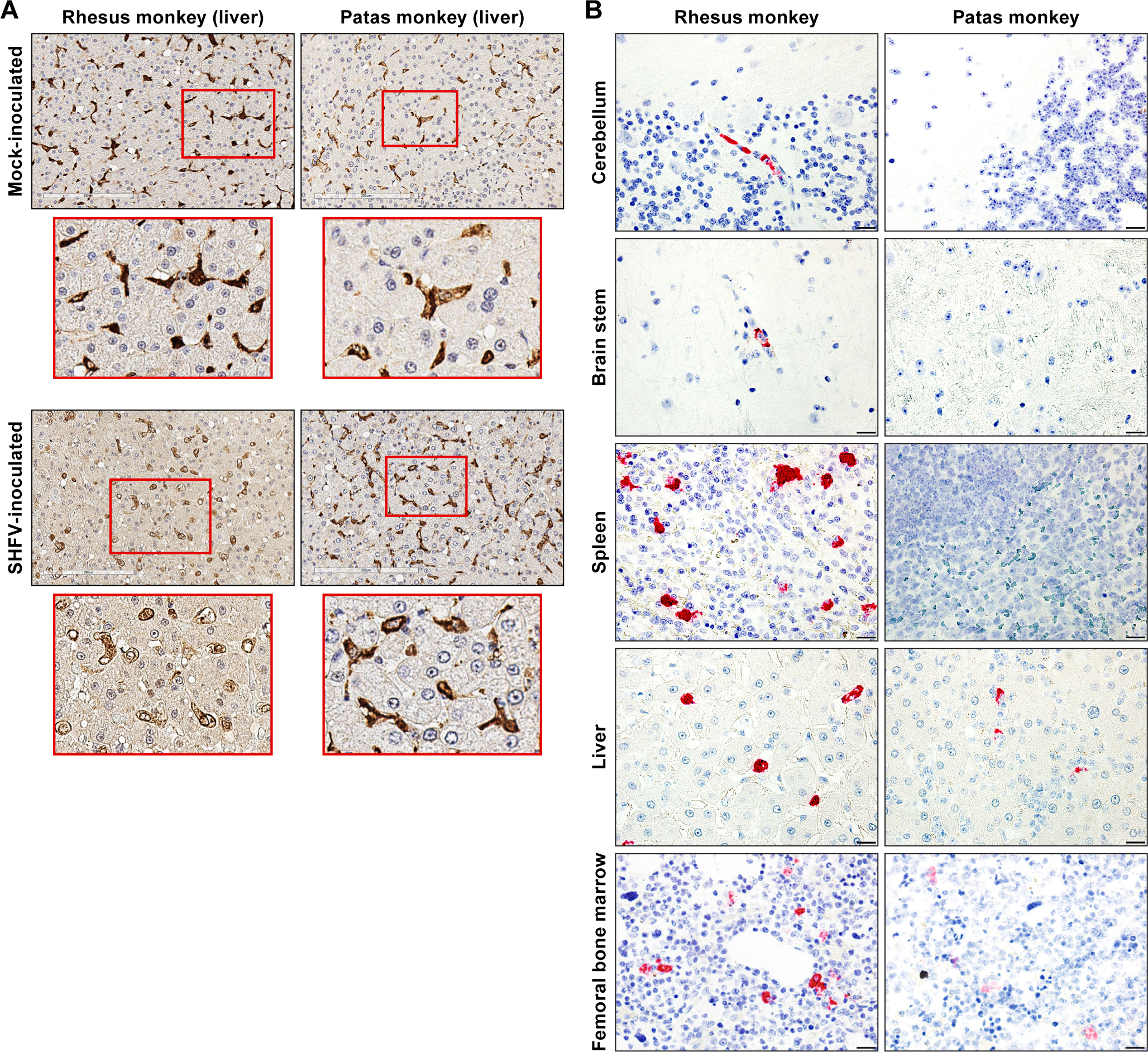
Representative images of liver immunohistochemistry for Iba1 in livers of mock- and SHFV-inoculated patas and rhesus monkeys; inset images highlight macrophage morphology for each group (A). Representative images of *in situ* hybridization for SHFV viral RNA in terminal cerebellum, brain stem, spleen, femoral bone marrow and liver samples from patas and rhesus monkeys inoculated with SHFV (B).

### Viremia is sustained in SHFV-infected patas monkeys

SHFV-inoculated patas and rhesus monkeys became viremic on day 2 PI (Fig 3A). The average peak titers in patas and rhesus monkeys were 6.75 (range 6.41–6.96) and 7.08 (range 6.79–7.36) log_10_ viral RNA (vRNA) copies per ml, respectively. Viremia peaked on day 4 PI in patas monkeys and was detectable in all three subjects for the remainder of the experiment, except for days 15 and 19 PI where viremia was below the limit of detection in two subjects. Terminal viremia in SHFV-inoculated patas monkeys was 3.42 (range 2.51– 4.43) log_10_ vRNA copies per ml. In SHFV-inoculated rhesus monkeys, viremia peaked between days 5 and 12 PI. Terminal viremia in SHFV-inoculated rhesus was 6.30, 7.36 and 4.17 log_10_ vRNA copies per mL in the two non-survivors and single survivor, respectively.

**Figure 3.**
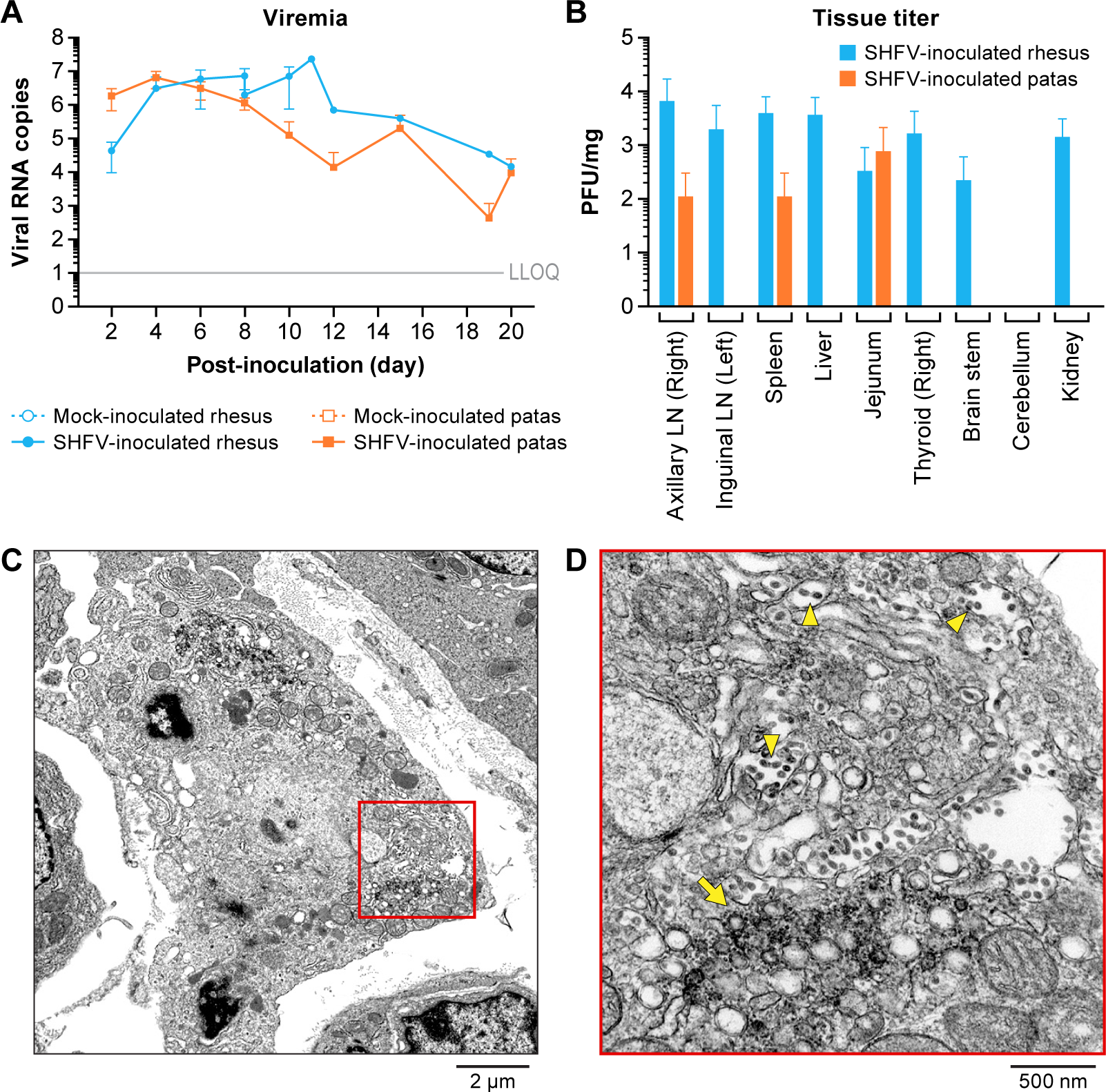
Mean viremia values in viral RNA copies per ml of whole blood for mock (solid symbols) and SHFV-inoculated (open symbols) patas monkeys (orange) and rhesus monkeys (blue) (A). Mean titer of tissues for SHFV-inoculated patas monkeys (orange) and rhesus monkeys (blue) in PFU per mg of 10% tissue homogenate (B, Lymph Node (LN)). Electron micrograph of jejunum from a SHFV-inoculated patas monkey showing double-membrane vesicles (C). Electron micrograph of jejunum from a SHFV-inoculated patas monkey showing apparently mature virions (yellow arrowheads) (D)

SHFV was detected by plaque assay in the axillary lymph node (n=1), spleen (n=1), and jejunum (n=1) of 2 SHFV-inoculated patas monkeys (Fig 3B). The highest titer was 3.36 log_10_ PFU/mg in the jejunum of the SHFV-inoculated patas with the highest terminal viremia. In the 2 non-surviving SHFV-inoculated rhesus monkeys, SHFV was found in axillary and inguinal lymph nodes (n=2 and 1, respectively), spleen (n=2), liver (n=2), jejunum (n=1), thyroid (n=2), brain-stem (n=1), and kidney (n=2). Only the kidney of the surviving SHFV-inoculated rhesus monkey contained SHFV, and its titer (2.53 log_10_ PFU/mg) was similar to those of the two non-survivors (2.82 and 5.35 log_10_PFU/mg, respectively). The highest tissue titer in SHFV-inoculated rhesus monkeys was 4.27 log_10_ PFU/mg in axillary lymph nodes.

Bone marrow, cerebella, jejuna, axial and inguinal lymph nodes, kidneys, and thyroids were assessed for evidence of SHFV infection by TEM. DMVs and apparently mature virions were found in the jejunum of the SHFV-inoculated patas monkey with the highest terminal viremia (Fig 3C and D). ISH for SHFV vRNA was performed to assess the livers, spleens, brainstems, cerebella, and femoral bone marrow of SHFV-inoculated subjects for signs of SHFV replication. The liver of 2 SHFV-inoculated patas monkeys and, rarely, the spleen of the third was positive for SHFV vRNA (Fig 2B) using RNAscope. The femoral bone marrow of the patas monkey with the highest terminal titer was positive for vRNA. vRNA was detected by ISH in the cerebellum, brain-stem, spleen, and liver of all SHFV-inoculated rhesus monkeys. vRNA was detected in femoral bone marrow of all three SHFV-inoculated rhesus monkeys. Morphologically, ISH data support that monocytes and endothelial cells are sites of SHFV infection in examined livers, spleens, brainstems, and cerebella of both SHFV-inoculated patas and rhesus monkeys. In all tissues, cells positive for SHFV vRNA (RNAScope) were morphologically consistent with macrophage-lineage cells; in each of the tissues evaluated these cells were present in fairly low numbers.

### SHFV infection of patas and rhesus monkeys elicit strong and overlapping immune responses

Quantitative immunohistochemistry (qIHC) revealed statistically significant (t-test, p<0.05) changes in inflammatory cell populations in livers of SHFV-inoculated monkeys (Fig 4A). On average, SHFV-inoculated patas monkeys had livers with increased CD3 and Iba1 signals when compared to uninfected patas monkeys. CD8 signals were increased in the liver of SHFV-inoculated patas monkeys when compared to uninfected patas monkeys but did not reach significance (p=0.06). CD8 and NKG2A signals were increased in SHFV-inoculated rhesus monkeys compared to uninfected controls. In the spleen, significant changes were seen in major histocompatibility complex class-1 (MHC1) and Iba1 signals between SHFV-inoculated and mock-inoculated patas monkeys (Fig 4B). No significant differences in cell concentrations were observed for CD3, CD8, and NKG2A in the splenic tissue of infected and uninfected patas monkeys. No significant changes were seen between SHFV-inoculated and mock-inoculated rhesus monkey spleens for any of the markers quantified.

**Figure 4.**
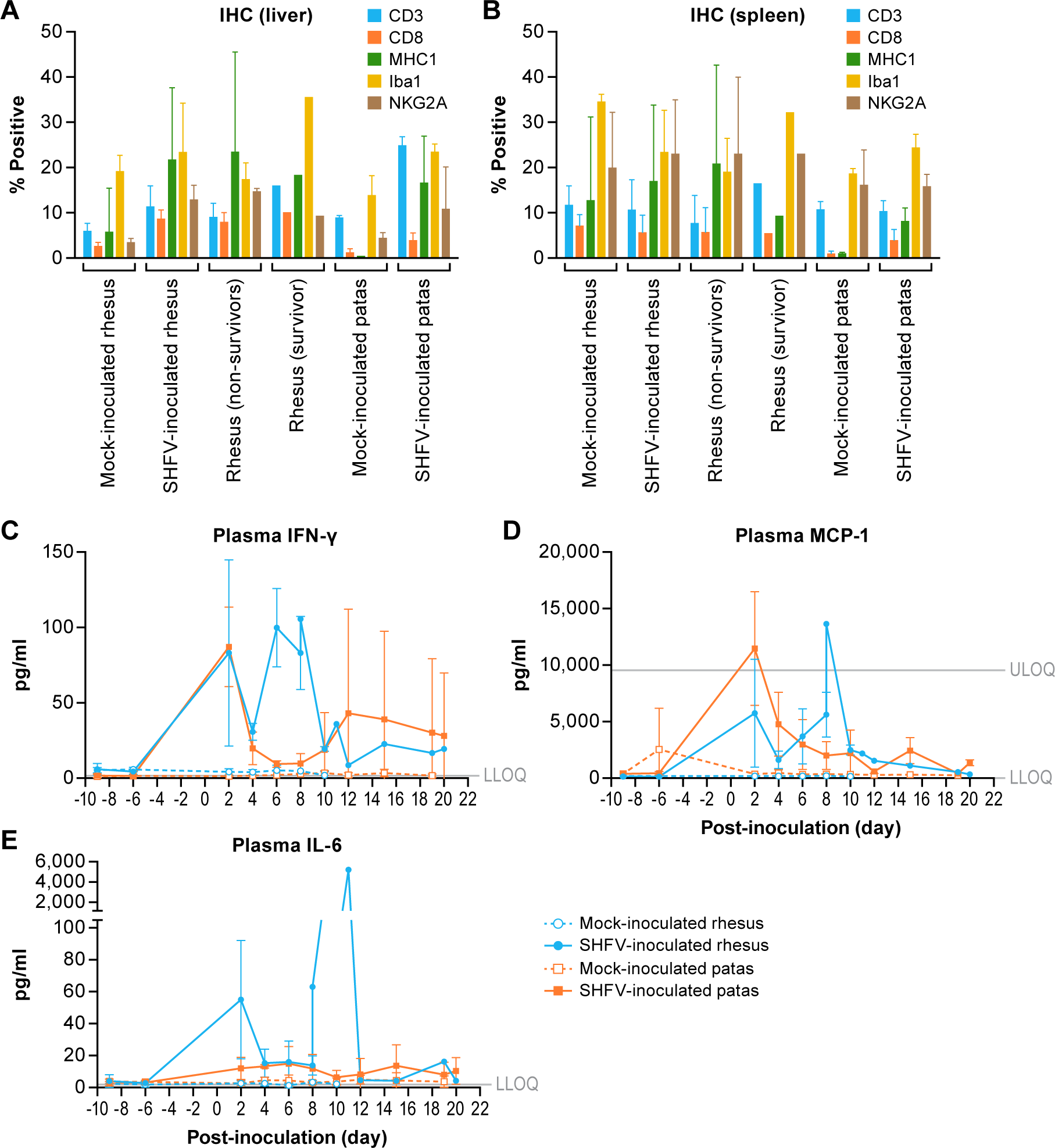
Mean quantitative immunohistochemistry values of indicated marker in mock- and SHFV-inoculated patas and rhesus monkey livers and spleens (A, B). Mean plasma concentrations in pg per ml of indicated analyte for mock (closed symbols) and SHFV-inoculated (open symbols) patas monkeys (orange) and rhesus monkeys (blue) (C–E). Gray lines represent the lower and upper limits of quantitation (LLOQ and ULOQ, respectively).

Interferon gamma (IFN-γ) concentrations in all SHFV-inoculated patas monkeys were elevated on day 2 PI compared to the pre-exposure mean concentration (group mean fold change (GMFC): 55.52, group mean concentration (GMC): 87.24 pg/ml) (Fig 4A). Later IFN-γ concentrations decreased to baseline in 2 SHFV-inoculated patas monkeys, whereas the concentration of the third patas monkey remained elevated throughout the experiment with a second peak in concentration at 12 days PI (group mean fold change (GMFC): 29.75; GMC: 43.11 pg/ml). The two non-surviving SHFV-inoculated rhesus monkeys had peak concentrations of similar magnitudes 2 days PI (GMFC: 14.57; GMC: 83.09 pg/ml) and all three subjects had increased concentrations by day 6 PI (GMFC: 20.58; GMC: 99.93 pg/ml).

Mean monocyte chemoattractant protein 1 (MCP-1) concentrations peaked at day 2 PI in all SHFV-inoculated patas monkeys and the two non-surviving rhesus monkeys (28.61 and 38.80 group MFC, respectively; 11,477.68 and 8,266.47 pg/ml GMC, respectively) (Fig 4B). All SHFV-inoculated rhesus monkeys had a second MCP-1 concentration peak at day 8 PI (group MFC: 50.68; GMC: 5,617.24 pg/ml).

SHFV-inoculated patas monkeys had mild increases in interleukin 6 (IL-6) concentrations in comparison to their mock counterparts (3.93 and 1.07 mean of individual PI fold changes respectively, 11.04 and 3.97 pg/ml GMC, respectively). SHFV-inoculated rhesus monkeys had increased IL-6 concentrations 2 days PI (group MFC: 17.34; GMC: 55.02 pg/ml) and the two non-survivors on their respective terminal days (419.82-fold change, 2,637.57 pg/ml non-survivor group mean concentration) (Fig 4E). The remaining analytes were below the limit of detection and were not considered for further analysis.

Flow cytometry of whole blood revealed increased numbers of circulating natural killer (NK) cells in SHFV-inoculated patas monkeys on days 12 and 19 PI (Fig 5A). In contrast, SHFV-inoculated rhesus monkeys had a single, larger, increase in circulating NK cells on day 8 PI. Changes in Ki67^+^ NK cells in SHFV-inoculated patas monkeys were more variable, with one patas monkey reaching peak numbers at 2 days PI and the other two patas monkeys reaching peak numbers of day 8 PI (Fig 5B). Increased Ki-67^+^ numbers in the surviving rhesus monkey returned to baseline by the conclusion of the experiment.

**Figure 5.**
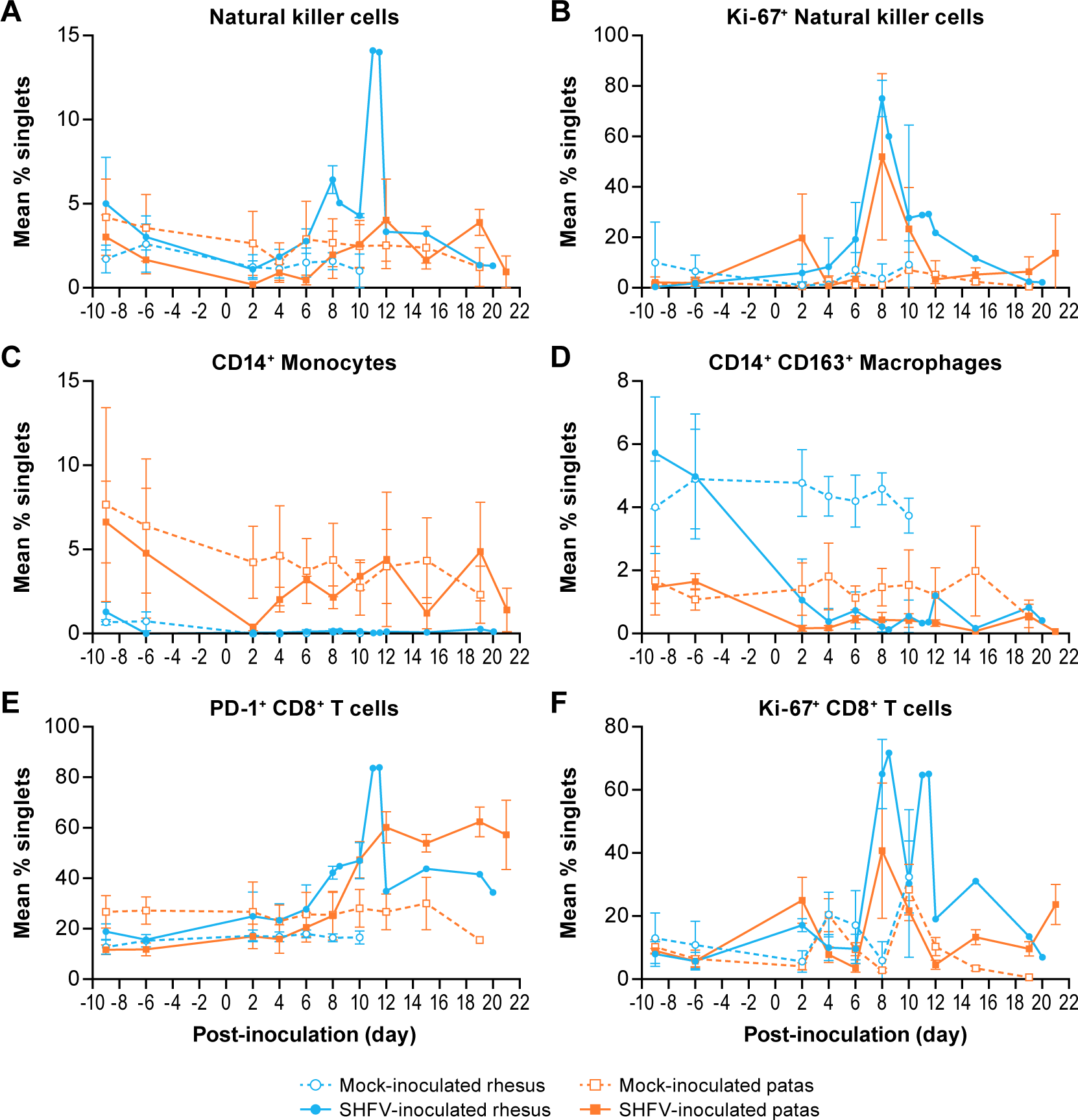
Mean percentage of indicated cell populations from whole blood of mock (closed symbols) and SHFV-inoculated (open symbols) patas monkeys (orange) and rhesus monkeys (blue) (A-F).

Circulating CD14^+^ monocytes were decreased in SHFV-inoculated patas monkeys at day 2 PI prior to returning to baseline counts, whereas counts in SHFV-inoculated rhesus monkeys appeared unchanged throughout the experiment (Fig 5C). SHFV-inoculated patas and rhesus monkeys had decreased numbers of CD14^+^ CD163^+^ macrophages (Fig 5D) starting at day 2 PI that remained until each subject’s respective endpoint. Both SHFV-inoculated patas and rhesus monkeys had declines in circulating CD4^+^ and CD8^+^ T-cell numbers starting on day 2 PI, but counts recovered by day 10 PI (data not shown). Numbers of PD-1^+^ CD8^+^ T-cells began to increase in both SHFV-inoculated patas and rhesus monkeys on day 10 and 8 PI, respectively, and remained elevated until the conclusion of the experiment (Fig 6E). Ki-67^+^ CD8^+^ T-cell numbers were elevated on day 2 PI before decreasing to baseline counts. A second, larger, peak in Ki-67^+^ CD8^+^ T-cell numbers was seen on day 8 PI in both SHFV-inoculated patas and rhesus monkeys, followed again by a return to baseline counts (Fig 5F).

**Figure 6.**
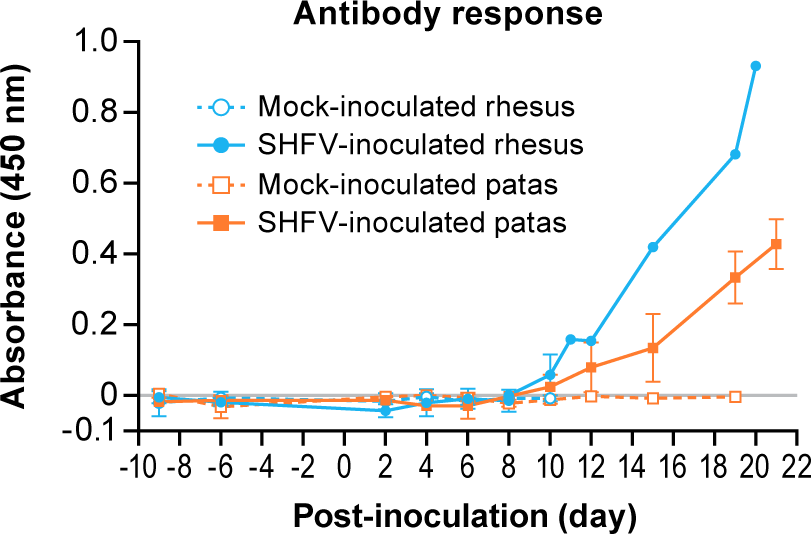
Mean ELISA absorbance values for patas monkeys (orange) and rhesus monkeys (blue) either inoculated with 5,000 PFU of SHFV (open symbols) or with PBS (closed symbols).

IgG antibody responses were detected by enzyme-linked immunosorbent assay (ELISA) in all three SHFV-inoculated patas monkeys and two of the three (the survivor and one non-survivor) SHFV-inoculated rhesus monkeys (Fig 6). Two SHFV-inoculated patas monkeys had detectable anti-SHFV antibodies on day 10 PI and the third on day 15 PI. The two SHFV-inoculated rhesus monkeys with a response began responding on day 10 PI. Response magnitude continued to increase in all responding subjects till their respective endpoints. Anti-SHFV antibodies were not detected in mock-inoculated subjects at any time.

## DISCUSSION

The goal of this study was to characterize SHFV infection of patas monkeys in comparison to rhesus monkeys to assess the usefulness of comparing biologically similar refractory and susceptible primates of different species in hemorrhagic fever virus infection. This is the first report of successful experimental SHFV infection of patas monkeys and further characterizes SHFV in rhesus monkeys.

Our data demonstrate that SHFV can replicate to high titers in patas monkeys without causing significant disease. Consistent with previously obtained data derived from experimentally SHFV-infected macaques (<u>10</u>, <u>13</u>, <u>14</u>), ISH and electron microscopy of infected tissues indicate that tissue-resident macrophages and endothelial cells are likely the main targets of SHFV in patas monkeys. Although not definitive, our findings indicate that SHFV may replicate in the same or similar cell populations in both patas and rhesus monkeys. Additional experiments are required to confirm the tissue distribution of SHFV and characterize the infected cell types over the course of the infection.

The immune response of SHFV-inoculated patas monkeys is similar to that of SHFV-inoculated rhesus monkeys and includes initial lymphopenia and monocytopenia, elevated IFN-γ and MCP-1 concentrations, and changes in circulating macrophage, NK, and T-cell populations. Patas monkeys did not react with IL-6 concentration increases to SHFV infection, whereas increased IL-6 concentrations in SHFV infected rhesus were observed in this experiment and have been previously reported (<u>10</u>, <u>13</u>, <u>14</u>). This difference is of particular interest as IL-6 has been associated with non-survival in SHFV-infected rhesus monkeys, and because decreased concentrations of IL-6 were seen in *in vitro* infection of monocyte-derived macrophages and dendritic cells from baboons (<u>13</u>, <u>15</u>). Given the potential role of IL-6 in human VHFs, our model offers an opportunity to explore the potential of therapies, such as neutralizing antibodies, aimed at modulating IL-6 responses during infection.

IFN-γ may be a more challenging target given that concentrations fluctuated in both patas and rhesus monkeys and that observed differences were largely temporal. Future experiments are required to determine whether the difference in IFN-γ responses between patas and rhesus monkeys is indicative of functional differences of NK or T-cells during SHFV infection. IFN-γ has been implicated in conflicting roles in other VHF-causing virus infections, such as filovirus and mammarenavirus infections, ranging from detrimental to protective (<u>21–26</u>). If differences exist between activation states of NK or T-cells between animals of different species, further characterization of relevant cytokine concentrations and cell-cell interactions is warranted. Given the temporal nature of the differences in IFN-γ, a more prudent therapeutic approach may be to directly stimulate or inhibit dysfunctional cell-types through supplementing factors or inhibiting signaling.

Livers of SHFV-inoculated patas monkeys had increased CD3 and Iba1 signals whereas those of SHFV-inoculated rhesus monkeys had elevated CD8 and NKG2A signals. Increases in CD8 and NKG2A signals in the absence of an increase in CD3 signal suggests that SHFV infection leads to an increase in infiltrating NK cells in rhesus monkeys (<u>27</u>). Indeed, the numbers of circulating NK cells were elevated at day 8 PI. Although there were clear quantitative differences in tissue and circulating NK cells between SHFV-infected patas and rhesus monkeys, all animals were affected by similar changes in circulating Ki-67^+^ NK cell numbers. The data suggest that a potent type 1 interferon response occurs in SHFV-inoculated rhesus monkeys, but it is unclear whether a similar response is present in SHFV-inoculated patas monkeys. NK cell responses are important for survival in human Ebola virus disease cases (<u>28</u>, <u>29</u>). Differences in the timing of NK cell responses are a key difference in non-lethal, mild disease and lethal, severe (Lassa virus) infections in macaques (<u>30</u>), suggesting the species-specific NK cell responses observed here warrant further exploration. For comparison, and as described in the accompanying paper by Buechler *et al*., olive baboons and rhesus monkeys infected with SWBV-1 also developed increases in NK and CD8^+^ T-cell numbers, with CD8^+^ T-cell numbers remaining elevated through the observation period. NK cell dynamics suggest a short-lived peak in olive baboons and rhesus monkeys that did not develop severe disease due to SWBV-1 infection. Together, these data support a role for appropriate timing and activation of NK cells in modulating disease presentation in VHFs.

Increased detection of Iba1, a macrophage marker, suggests that SHFV infection leads to an increase in the number of macrophages in the liver (<u>31</u>). Determining the sources of these additional liver macrophages in SHFV-inoculated patas monkeys is important given the changes in MCP-1 concentrations and the differential roles of hepatic resident and non-resident macrophages (<u>32–35</u>). SHVF is dependent on CD163 for cellular entry (<u>36</u>), and preferential targeting of CD163^+^ cells by SHFV may explain why both patas and rhesus monkeys lost CD163^+^ macrophages. CD163 is associated with alternatively-polarized macrophages and macrophage polarization has a significant role in immunity and infection (<u>37–39</u>). Additionally, the presence of rounded and vacuolated macrophages in the livers of SHFV-inoculated rhesus monkeys but not their patas counter parts suggests and active response to infection and that tissue-resident macrophages may play a key role in SHFV pathogenesis. Future serial-sampling studies will be required to fully characterize tissue-specific immune responses over the course of SHFV infection, but will be challenging since the disease is not uniformly lethal.

Consistent with the complete blood counts, flow cytometry revealed early decreases in both CD4^+^ and CD8^+^ T-cells supporting the presence of lymphopenia in both patas and rhesus monkeys. Interestingly, on day 2 PI, both SHFV-inoculated patas and rhesus monkeys had an increase in Ki-67^+^ CD8^+^ T-cell numbers, suggesting some level of proliferation was present even in the face of depletion (<u>40</u>, <u>41</u>). On days 8 and 10 PI, CD8^+^ T-cell counts returned to baseline in all monkeys. This return coincided with a second spike in Ki-67^+^ CD8^+^ counts and the start of an elevated count of PD-1^+^ CD8^+^ T-cells. PD-1 is associated with T-cell exhaustion and is also found on activated T-cells (<u>42</u>, <u>43</u>). PD-1^+^ T-cells and CD8^+^ T-cells appear to play important roles in human cases of Ebola virus disease (<u>28</u>, <u>29</u>, <u>44–46</u>). Consequently, it may be fruitful to determine if SHFV provides a means for studying the role of T-cell dysfunction in EVD. As SHFV is capable of infecting antigen-presenting cells, it will be important to determine if poor T-cell responses during SHFV infection are due to deficits in antigen presentation as is suspected to be the case during Ebola virus infection in humans and non-human primates (<u>21</u>, <u>46–50</u>). T-cell exhaustion has also been implicated in chronic viral infections and may be an important mechanism in the development of persistent SHFV infections in patas monkeys (<u>43</u>). Given the extreme differences in disease course and outcomes during SHFV infection in patas and rhesus monkeys the relatively high degree of overlap in host-response features was unexpected. Changes in IFN-γ and MCP-1 concentrations, and circulating macrophage, NK, and T-cell populations were similar between SHFV-inoculated patas and rhesus monkeys. Interestingly, viral loads were largely similar between all monkeys and ISH data suggests that SHFV targets the same cell types in cercopithecines. Our data suggests that when biologically similar primates of two species are infected with the same pathogen that host responses are initially quite similar however, as the virus interacts with the host to modulate the immune response, the effectiveness of the immune response is diminished, likely as a result of the degree of adaptation to the host.

Rather than a “cytokine storm”, we propose that a process driven by NK cells and macrophages is the deciding factor in developing hemorrhagic fever, either alone, or through their impacts on otherwise beneficial or seemingly inconsequential host-responses. Ultimately, the differences in disease course in SHFV patas and rhesus monkeys are due to their biological differences, ranging from organism-level physiology to minute genotypic differences. Although biologically similar, patas and rhesus monkeys are distinct species. A core drawback of the approach taken in the paper is that although it is relatively simple to catalogue host-responses in monkeys of two species, it is difficult to contextualize them. Without extensive characterization, it is challenging to determine which impact a given host-response has on members of each species: clear differences may have no impact on disease whereas topically similar responses may have divergent functional impacts.

However, as a tool for identifying host-response mechanisms as targets for medical countermeasures against viral hemorrhagic fevers, the issue of biological differences may become more tractable. Interactions between cell-cell and cell-SHFV interactions may be driven by highly-specific mechanics and are therefore far more difficult to translate or target. On the other hand, the resulting functional differences are more readily targetable. For example, it may be that the large differences in IL-6 concentrations between patas and rhesus monkeys observed in this work are driven by SHFV-infected macrophages. Rather than attempting to navigate the convolutions of species-specific macrophage biology, targeting IL-6-producing macrophages, IL-6, or the IL-6 receptor offer far more direct paths to identifying druggable targets. Under the macrophage/IL-6 example and in the context of identifying targets for therapeutic development, the minutiae of species-specific macrophage biology may be rather unimportant when the results of those biologies suggest a common target.

A top down approach to more fully characterize SHFV infection in patas and rhesus monkeys will streamline identification of host components or sub-systems that are sources of responses that result in VHF. Identification of and targeting the high-level responses could ultimately even result in pan-VHF therapeutics.

## MATERIALS AND METHODS

### Cells and Virus

Simian hemorrhagic fever virus (SHFV; *Nidovirales: Arteriviridae: Simartevirus: Simian hemorrhagic fever virus*) strain LVR42-0/M6941 (<u>51</u>) was passaged twice before a final passage on MA-104 cells to create the virus stock. Briefly, virus stock was prepared by freeze-thawing infected cells three times prior to clarification with low-speed centrifugation and was then concentrated by centrifugation at 16,000xg. Pellets were resuspended in PBS and combined. The final viral stock was sequenced as in (<u>52</u>) for quality control purposes (GenBank accession number pending). Sequencing confirmed the expected genotype and lack of any contamination.

### Animals

Six patas monkeys (4 females and 2 males) and 6 rhesus monkeys (3 females and 3 males) where used in this study. Patas monkeys ranged from 5.51–14.01 kg in weight and 9–14 years in age, whereas rhesus monkeys ranged from 4.77–12.75 kg in weight and 8–12 years in age. Rhesus monkeys were obtained from the National Institute of Allergy and Infectious Disease (NIH/NIAID) rhesus monkey colony. Patas monkeys were obtained from the National Institute of Child Health and Human Development (NIH/NICHD). Subjects were screened for simian T-lymphotrophic virus, simian immunodeficiency virus, and simian retrovirus infections, and cleared for use in the experiment by the facility veterinarian. Patas monkeys were determined to be serologically negative for SHFV prior to enrollment. Rhesus monkeys were obtained from a SHFV-free source, and therefore were not screened prior to use in this experiment. Subjects were randomly assigned to 4 groups (2 groups of patas monkeys, inoculated and mock-inoculated, and 2 groups of rhesus monkeys, inoculated and mock-inoculated), for sex, age and weight. The animals of one group of patas monkeys and 1 group of rhesus monkeys each received 5,000 PFU of SHFV diluted in 1 ml of PBS, whereas the animals of the remaining groups each received 1 ml of PBS (mock) by intramuscular injection of the right quadriceps. Subjects were housed in separate rooms and had access to food and water *ad libitum*.

Subjects were monitored at least twice daily. Physical exams were performed on pre-determined experimental days (−9, −6, 0, 2, 4, 6, 8, 10, 12, 15, 19, and 21) and prior to euthanasia. Blood was collected on all physical exam days, except day 0, and prior to euthanasia. Scheduled days for euthanasia with necropsy were as follows: mock-inoculated patas monkeys at 19 days post-inoculation, SHFV-inoculated patas monkeys at 21 days post-inoculation, mock-inoculated rhesus monkeys at 10 days post-inoculation, and SHFV-inoculated rhesus monkeys at 20 days post-inoculation. Subjects were euthanized at scheduled times or upon reaching pre-established clinical endpoint criteria including overall clinical appearance, respiratory function, responsiveness, and core body temperature. At euthanasia, subjects were perfused with saline before necropsy and sample collection. Subjects were housed in an AAALAC, International, accredited facility under biosafety level 4 (BSL-4) conditions due to the nature of the facility in which the experiments were performed. All experimental procedures were approved by the NIAID DCR Animal Care and Use Committee and were performed in compliance with the Animal Welfare Act regulations, Public Health Service policy, and the Guide for the Care and Use of Laboratory Animals recommendations.

### Virus Quantification

Virus stock and tissue concentrations were determined by plaque assay on MA-104 cells. Briefly, serial dilutions of 10% (w/v) tissue homogenates were added to cell monolayers and incubated for 1 h. Then, monolayers were overlaid with 0.8% tragacanth (Sigma, St. Louse, MO), MEM (Lonza, Walkersville, MD), 1% penicillin-streptomycin (Lonza) and 2% heat-inactivated fetal bovine serum (Sigma) final concentration. After a 3-day incubation, overlays were aspirated, and monolayers fixed using 10% neutral-buffered formalin (Fisher Scientific, Hampton, TN) with 0.2% crystal violet (Ricca, Arlington, TC) prior to enumeration.

### Plasma Cytokines

Plasma concentrations of granulocyte-macrophage colony stimulating factor (GM-CSF), interferon gamma (IFN-γ), macrophage chemoattractant protein 1 (MCP-1), vascular endothelium growth factor (VEGF), and interleukins 2, 4, 6, 8, 10, and 17 were measured using a Milliplex non-human primate kit (MilliporeSigma, St Louis, MO) as described previously (<u>13</u>).

### Hematology

Complete blood counts (Sysmex XS1000i, Lincolnshire, IL) and selected serum chemistries using Piccolo General Chemistry 13 kits (Abaxis, Union City, CA) were performed at the described timepoints using blood collected in K3 EDTA and SST tubes (BD, San Jose, CA). Due to a lack of published data, standard ranges for patas monkeys were defined as the mean +/− two standard deviations of all pre-exposure timepoints. Standard ranges for rhesus monkeys were determined from data kept by veterinary staff on subjects housed in the facility.

### Histology, *In Situ* Hybridization, and Immunohistochemistry

Formalin-fixed paraffin-embedded (FFPE) animal tissue sections (5 µm) were used for immunohistochemical staining using the following antibodies: NKG2A (Abcam, Cambridge, MA); Iba1 (Wako, Richmond, VA); MHC1 [Clone EPR1394Y] (Abcam); CD8 (Abcam); CD3 [Clone 12] (AbD, Serotec Hercules, CA). Staining was performed on the Bond RX platform (Leica Biosystems) according to the manufacturer’s protocol. Briefly, sections were baked, deparaffinized, and rehydrated. Epitope retrieval was performed using Leica Epitope Retrieval Solution 1, pH 6.0, heated to 100°C for 20 min, and quenched with hydrogen peroxide prior to addition of primary antibody. The Bond Polymer Refine Detection kit (Leica Biosystems) was used for chromogen detection. Image analysis was performed on select tissues from all groups to quantify the degree of positive staining. Images were obtained on a bright-field Leica Aperio AT2 slide scanner (Leica Biosystems) and processed using Aperio Image Scope [v12.3] algorithms. The Positive Pixel Count Algorithm was used to assign pixels to intensity ranges for positive (strong (n_sp_), medium (n_p_), and weak (n_wp_)) and negative (n_n_) pixels. Pixels were categorized, and positive percentage was calculated per image as a fraction of the number of strong positive (n_sp_) pixel to the total number of stained pixels:

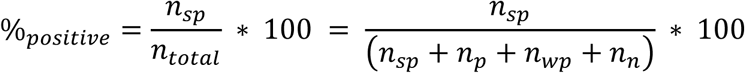

SHFV RNA RNAscope *in situ* hybridization (ISH) was performed as previously described (<u>53</u>).

### Electron Microscopy

Electron microscopy samples were processed and imaged as previously reported (<u>54</u>).

### qPCR viremia

Monkey peripheral blood samples were inactivated in 3 volumes of Trizol LS buffer (Thermo Fisher Scientific, Waltham, MA) and RNA was extracted using the Qiagen AllPrep 96 kit as described by manufacturer (Qiagen, Valencia, CA) except that each sample was treated with 27 units of DNAse I (Qiagen). SHFV RNA copy number was determined by RT-qPCR using primers and probes targeting the SHFV gp15 nucleocapsid gene. The AgPath-ID One-Step RT-PCR kit (Thermo Fisher Scientific) was used to perform the assay. Primers and Cal Fluor Orange 560/BHQ1 labeled probe were synthetized by LCG Biosearch Technologies (Petaluma, CA). RT-qPCR reactions were performed in 20-µl reactions using forward primer (5′-CGACCTCCGAGTTGTTCTACCT-3′), reverse primer (5′-GCCTCCGTTGTCGTAGTACCT-3′), and fluorescent probe (5′-CCCACCTCAGCACACATCAAACAGCT-3′). Synthetic DNA (5′-TTTCGCCGAACCCGGCGACCTCCGAGTTGTTCTACCTGGTCCCACCTCAGCACAC ATCAAACAGCTGCTGATCAGGTACTACGACAACGGAGGCGGAAATCTTTCATATG-3′; LCG Biosearch Technologies, Novato, CA) was used as a standard. The qPCR reactions were run at 50 °C for 10 min, 95 °C for 10 min, 55 cycles of 95 °C for 15 s, and 60 °C for 45 s on a 7900HT Fast Real-Time PCR System (Thermo Fisher Scientific). Data were analyzed using Applied Biosystems 7900HT Fast Real-Time PCR System Software (Thermo Fisher Scientific).

### Flow Cytometry

Whole blood was assessed for the following markers: HLA-DR-FITC (BioLegend, San Diego, CA), CD16-APC (BD), PD-1-APC-Cy7 (BioLegend), CD3-AF700 (BD), CD11c-PE (BD), CD28-PE-Cy5 (BioLegend), NKG2a-PE-Cy7 (Beckman Coulter, Brea CA), CD163-PE/Dazzle594 (BioLegend), CD4-BV421 (BD), CD14-BV510 (BioLegend), CD20-BV570 (BioLegend), CD8a-BV605 (BD), CD123-BV650 (BD), PD-L1-BV711 (BioLegend), CD95-BV785 (BioLegend), and Ki-67-PerCP-Cy5.5 (BD). Briefly, 100 μl of whole blood was incubated with 100 μl of the marker panel (excluding Ki-67) and incubated for 20 min. Red blood cells were lysed with 1 ml of BD FACSLyse (BD) for 10 min. Cells were washed and fixed for 30 min using BD Cytofix/Cytoperm (BD) and incubated with Ki-67 antibody for 30 min at 4° C, followed by a final wash in 1xBD Permwash (BD). Data acquisition and analysis was performed with FlowJo version 10.2 (BD).

### Enzyme-linked immunosorbent Assays

Lysates from MA-104 cells infected with SHVF or mock infected (media only) were used as substrates. Immulon 2 HB microplates (Thermo Fisher Scientific, Walkersville, MD) were coated with cell lysates diluted in PBS and incubated overnight at 4◦C. Plates were then washed five times with wash buffer comprised of PBS/0.2% Tween 20 and blocked for 2 h at room temperature with 5% nonfat milk (LabScientific, Highlands, NJ, USA) dissolved in PBS. Plates were then washed five times with wash buffer, and analyte serum diluted at 1:50 in PBS/2.5% milk/0.05% Tween 20 was added in duplicate to corresponding wells. After a 1-h incubation at room temperature, plates were washed and horseradish peroxidase-conjugated anti-monkey IgG (Sigma Aldrich, St. Louis, MO, USA; A2054) was added. Plates were then incubated for 1 h at room temperature before washing with wash buffer and adding TMB Substrate (Thermo Fisher Scientific, Walkersville, MD, USA). Following a 10-min incubation, 100 μl of TMB stop solution (Thermo Fisher Scientific, Walkersville, MD, USA) were added to each well and the plates were read on a SpectraMax Plus 384 plate reader (Molecular Devices, Sunnyvale, CA, USA) at 450 nm.

## Acknowledgments

This work, in part, was supported by the NIAID Division of Intramural Research and the NIAID Division of Clinical Research via the Battelle Memorial Institute’s prime contract with the National Institute of Allergy and Infectious Diseases (NIAID) at the National Institutes of Health, under contract no. HHSN272200700016I (DP, KRHP, SM, JGB, JHK.

We would like to thank the entire EVPS and IRF-Frederick staff for their support of the experiments. We especially would like to thank Tim Cooper (IRF-Frederick) for his help in establishing standard ranges for complete blood counts and serum chemistries. We also would like to thank Laura Bollinger and Jiro Wada (IRF-Frederick) for technical writing services and figure preparation, respectively.

